# Frugal alignment-free identification of FLT3-internal tandem duplications with FiLT3r

**DOI:** 10.1101/2022.06.27.496265

**Authors:** Augustin Boudry, Sasha Darmon, Nicolas Duployez, Martin Figeac, Sandrine Geffroy, Maxime Bucci, Karine Celli-Lebras, Matthieu Duchmann, Romane Joudinaud, Laurène Fenwarth, Olivier Nibourel, Laure Goursaud, Raphael Itzykson, Hervé Dombret, Mathilde Hunault, Claude Preudhomme, Mikaël Salson

## Abstract

**Background:** Internal tandem duplications in the *FLT3* gene, termed *FLT3*-ITDs, are useful molecular markers in acute myeloid leukemia (AML) for patient risk stratification and follow-up. *FLT3*-ITDs are increasingly screened through high-throughput sequencing (HTS) raising the need for robust and efficient algorithms. We developed a new algorithm, which performs no alignment and uses little resources, to identify and quantify *FLT3*-ITDs in HTS data.

**Results:** Our algorithm (FiLT3r) focuses on the *k*-mers from reads covering *FLT3* exons 14 and 15. We show that those *k*-mers bring enough information to accurately detect, determine the length and quantify *FLT3*-ITD duplications. We compare the performances of FiLT3r to state-of-the-art alternatives and to fragment analysis, the gold standard method, on a cohort of 185 AML patients sequenced with capture-based HTS. On this dataset FiLT3r is more precise (no false positive nor false negative) than the other software evaluated. We also assess the software on public RNA-Seq data, which confirms the previous results and shows that FiLT3r requires little resources compared to other software.

**Conclusion:** FiLT3r is a free software available at https://gitlab.univ-lille.fr/filt3r/filt3r. The repository also contains a Snakefile to reproduce our experiments. We show that FiLT3r detects FLT3-ITDs better than other software while using less memory and time.

## 1 Background

Monitoring the disease progression in blood cancers requires the identification of pathology-specific molecular markers. In acute myeloid leukemia (AML), one of the strongest risk factor used for risk stratification is the presence of an internal tandem duplication in the *FLT3* gene (*FLT3*-ITD), which occurs in 20-30% of the cases [1],[2],[3],[4]. *FLT3*-ITD are in-frame duplications of highly variable size, ranging from 3 to more than 400 nucleotides, mostly located within the receptor’s autoinhibitory juxatamembrane domain. This mutation represents a strong prognostic biomarker since patients with *FLT3*-ITD are at higher risk of relapse and have decreased event-free and overall survival [5]. The importance of allelic ratio (AR), as assessed by the ratio between the mutated allele and the wild-type allele, has been demonstrated in several studies [6],[4]. Presence of *FLT3*-ITD has a major therapeutic impact, such as indication of allogeneic hematopoietic stem cell transplantation (allo-HSCT) in first complete remission [7],[8],[6]. *FLT3*-ITD is also a therapeutic target with the emergence of *FLT3* inhibitors, used in combination with chemotherapy during induction and consolidation courses [9],[10] and more recently as maintenance therapy [11].

Historically, according to European LeukemiaNet (ELN) guidelines, the identification and quantification of *FLT3*-ITD were performed with fragment analysis [7]. DNA fragments were fluorescently labeled, then separated by capillary electrophoresis [12],[13]. One or more peaks were obtained depending on the presence or absence of ITD(s). The size of the ITD was determined by subtracting the size of the wild type fragment from that of the mutated fragment, using a scale. The AR was evaluated by dividing the area under the curve of the mutated fragment’s peak by that of the wild-type fragment’s peak. Although robust, this technique has several limitations: (i) the lower limit of quantification of the AR is high, generally at least 1% [14],[15], (ii) the determination of the size of the ITD using the scale is approximate, (iii) the exact position of insertion and complete sequence of the ITD are not available [16],[17] and (iv) sample multiplexing cannot be performed (v) quantification of very large insertions might be biased. Development of high-throughput sequencing enabled the detection of several genetic alterations in a single sequencing run and has the potential to overcome several of those limitations. However, classical mappers, such as Bowtie2 or BWA, cannot directly identify structural variants such as tandem duplications. Thus initially, the data produced were analysed using pindel, a general-purpose algorithm that detects and quantifies indels and structural variants [18]. Although using a general-purpose software can be appealing, it may be under-optimised to address the specific problem of identifying and quantifying tandem duplications, especially *FLT3*-ITD. Thus, many methods were specifically developed to detect *FLT3*-ITD in high-throughput sequencing data [19]. Yuan and colleagues distinguished the assembly-based methods from the alignment-based ones, and demonstrated better accuracy of alignment-based methods.

However, alignment-free methods have gained importance in bioinformatics [20] as they usually have the advantage of using only a fraction of the resources required by alignment methods while providing similar results. km is an example of an alignment-free strategy for *FLT3*-ITD detection [21]. It is based on Jellyfish [22], a *k*-mer counting algorithm, to efficiently count *k*-mers in the reads. Then km first builds a linear *k*-mer graph of the reference sequence and traverses it using their count table. Any divergent path that can be identified with the count table is a potential duplication. We introduce a new alignment-free approach based on *k*-mers, which is faster than km because it doesn’t require counting the *k*-mers from the reads. For the detection of duplications, we were inspired by the methodology used in the work of [23] by analysing the occurrences of the *k*-mers in the reads.

## 2 Methods

We introduce our heuristic aimed at finding duplications compared to a reference sequence, though it can more generally detect insertions and deletions. In what follows what is described for duplication also holds for indels without loss of generality.

To determine if a read contains a duplication, we first need to consider all *k*-mers from the read and identify their positions in the reference sequence. The main principle consists of identifying any read in which *k*-mer positions on the reference sequence would suddenly go back. Such an event would correspond to a duplication with respect to the reference sequence. To further explain this principle, here is a concrete example of its application. Let *T* be the reference sequence and *R* a read. Fig. 1 illustrates an example where AGAT is duplicated in *R*. When querying at which positions the *k*-mers of *R* occur in *T*, the positions of the *k*-mers gradually increase from 3 to 5 and then go back to 4 at the start of the duplication. This phenomenon is the signature of a duplication. We use this observation to detect duplications efficiently. As will be shown later, this information is also sufficient to determine the length of the duplication.

**Figure 1.**
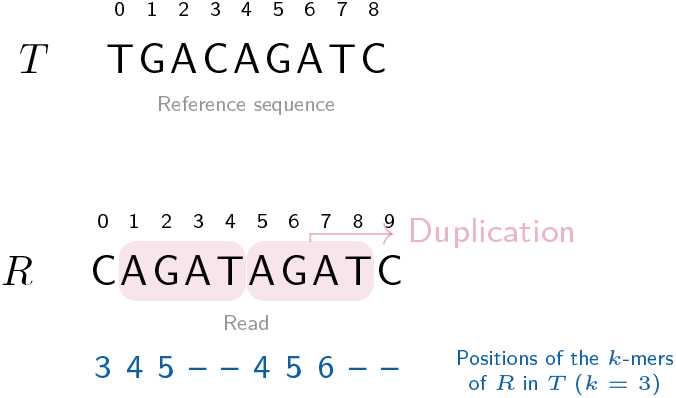
Identifying duplications from a reference sequence *T* in a read *R* with *k*-mers (*k* = 3). The bottom line represents the positions of the *k*-mers from *R* in *T*. For instance, the first element (3) corresponds to the *k*-mer CAG which appears in position 3 in *T*. The dashes (–) correspond to *k*-mers from *R* that do not occur in *T* (*eg*. ATA or TAG).

### 2.1 The FiLT3r algorithm

Our algorithm can be summarised in three main steps:

1. Indexing *k*-mers of the reference sequence with their original positions.
2. Traversing all the reads: keep the ones with enough *k*-mers from the reference sequence and determine the position of the read’s *k*-mers in the reference sequence.
3. Detect a duplication event in reads using the *k*-mers positions in the reference sequence.

The indexing step is trivial, as the reference sequence (basically exons 14 to 15 in the *FLT3* gene) is very short. With short values of *k*, there may exists several occurrences of the same *k*-mer in the reference sequence (see A in fig. 2). Thus, all positions are stored to prevent loss of information. Ambiguities in the positions will be resolved afterwards.

**Figure 2.**
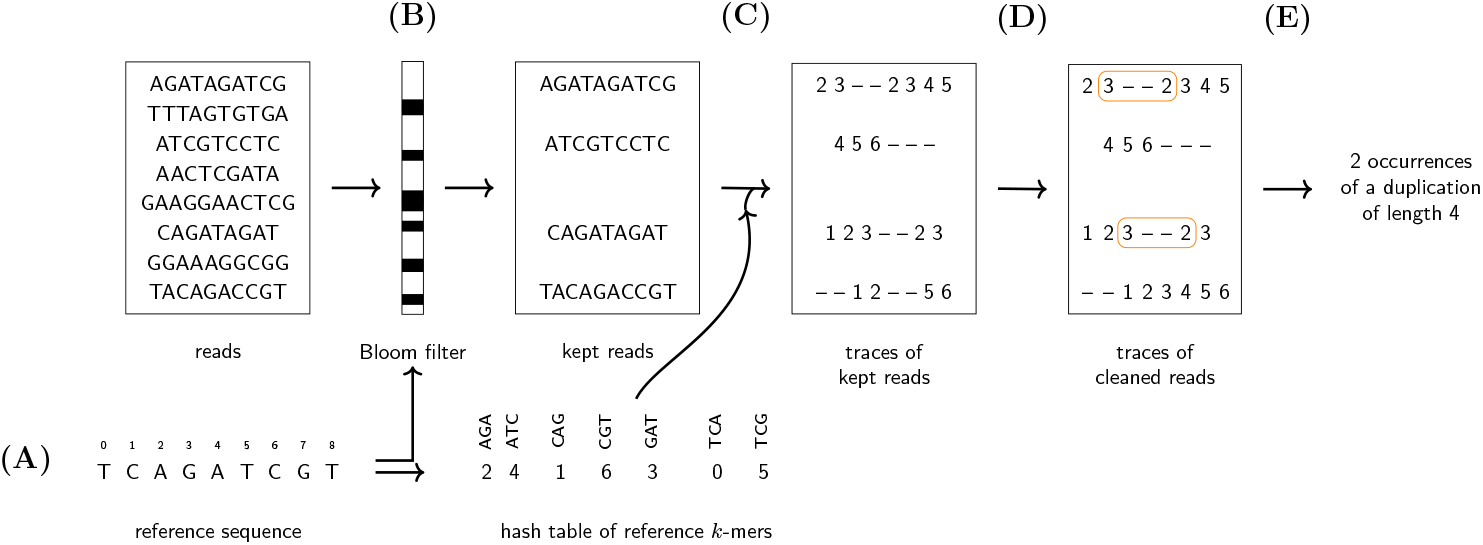
How FiLT3r processes the reads. (A) The *k*-mers of the reference sequence are indexed in a hash table and a Bloom filter. The keys of the hash table are *k*-mers and the values are a list of positions where the key occurs in the reference sequence. (B) The reads matching a sufficient number of *k*-mers with the reference are kept. (C) The *k*-mer positions in the reference are considered. (D) Substitutions are removed. (E) Duplications, and more generally indels, are called only using the positions of the *k*-mers.

The second step only consists of reading the reads one by one and then querying the index (a hash table) with the *k*-mers of each read (see C in fig. 2). The main difficulty comes from the third step. Fig. 1 introduces a simplified version of the problem, but other cases may arise that should be handled correctly to prevent false negatives or false positives as will be shown later.

Duplications are only searched in reads whose number of *k*-mers from the reference sequence is above a given threshold to prevent spending time on reads not coming from the gene of interest. For each of these reads we will now focus on the *k*-mers occurrences in the reference sequence. We call *trace of R in T*, denoted by *t*_*R*_, the list of *k*-mers positions in *T* from a read *R*. Then *t*_*R*_[*i*] is a list of positions where the *k*-mer *R*[*i* … *i*+*k* − 1] occurs in *T* (see *traces of kept reads* in fig. 2 for examples). This list can obviously be empty whenever the *k*-mer doesn’t occur in *T*. We call a break (as in [23]) the maximal span of positions where *t*_*R*_[*i*] is empty (denoted by *t*_*R*_[*i*] = –). Thus, a break is identified by its starting position and its ending position. For a read *R* if we have a break (*j*_1_, *j*_2_), then *trace*[*j*] = –, for each *j*_1_ ≤ *j* ≤ *j*_2_, and either *j*_1_ = 0 or *t*_*R*_[*j*_1_ − 1] ≠ –, and either *j*_2_ = |*R*| − 1 or *t*_*R*_[*j*_2_ + 1] ≠ –.

Whenever a duplication occurs in a read *R*, this will create a break in the trace. Note that the converse is not true: many other events can also create breaks. Actually, any difference between the read and the reference sequence will lead to a break. Also, in some rare cases, the duplication will not create a break. However, the probability of this phenomenon decreases exponentially with *k*.

Using the trace of the reads we start by identifying all the breaks in a trace and by determining if the positions before and after the break are compatible with a duplication. Thus, assuming for now that *t*_*R*_[*j*_1_ − 1] and *t*_*R*_[*j*_2_ + 1] both correspond to a single position in the reference sequence, we check whether *t*_*R*_[*j*_1_ − 1] + *g* − 1 > *t*_*R*_[*j*_2_ + 1], where *g* is the size of the break. In such a case, a duplication has been detected (see E in fig. 2).

The length of the duplication can be deduced from the *k*-mer positions. Let (*j*_1_, *j*_2_) be a break corresponding to a duplication. *t*_*R*_[*j*_1_ − 1] + *k* − 1 is the last nucleotide position before the duplication, while *t*_*R*_[*j*_2_ + 1] is the first nucleotide in the duplication. Thus the length of the duplication is *t*_*R*_[*j*_1_ − 1] + *k* − 1 − *t*_*R*_[*j*_2_ + 1] + 1 = *t*_*R*_[*j*_1_ − 1] −*t*_*R*_[*j*_2_ + 1] +*k*. For instance, in fig. 1, the break is (3, 4) and the length of the duplication is 5 − 4 + 3 = 4. However, the formula does not hold whenever short insertions occur at the breakpoint or when there is an overlap between the start of the duplication and what follows it in the reference sequence. All those cases can actually be easily dealt with. Each inserted nucleotide will lead to a break length increased by one. Conversely, each overlap will lead to a decreased gap length. We note that when no such insertion or overlap occurs, we have *j*_2_ − *j*_1_ + 2 = *k*. Hence, we deduce that more generally the duplication size is *t*_*R*_[*j*_1_ −1]−*t*_*R*_[*j*_2_ +1]+*j*_2_ −*j*_1_+2.

The formula also holds when substitutions occur at *d* < *k* nucleotides from the break-point (see fig. 3). In this case, this will make the break longer by *d* nucleotides, but this will also decrease *t*_*R*_[*j*_1_ −1] by *d* or increase *t*_*R*_[*j*_2_ +1] by *d* (depending where the substitution occurs). Thus, the computed duplication size won’t change compared to the situation where no such substitution occurs. Thus, our heuristic is robust to such events.

**Figure 3.**
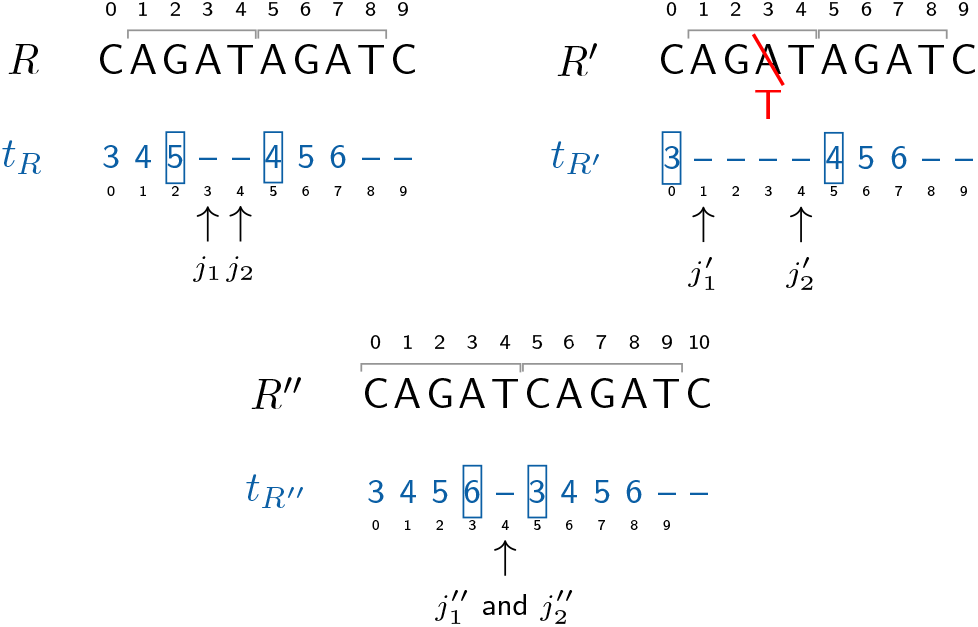
Computing the duplication length. Three examples are shown using the same reference as in fig. 1. In the first example (left) we also have the same read as in fig. 1. The break (*j*_1_, *j*_2_) = (3, 4) is due to a duplication. Its length will be computed with *t*_*R*_[*j*_1_ − 1] − *t*_*R*_[*j*_2_ + 1] + *j*_2_ − *j*_1_ + 2 = 5 − 4 + 4 − 3 + 2 = 4. In the second example (right) a single nucleotide is modified leading to a longer break (1, 4). Thus 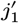 decreased by 2 compared to the first example. However, in the meantime 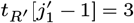 also decreased by 2 compared to the first example (where *t*_*R*_[*j*_1_ − 1] = 5). Thus, the duplication length is identical: 3 − 4 + 4 − 1 + 2 = 4. This is an example where our algorithm can detect the duplication even when it contains a substitution. The process is similar with an indel. In the third example (bottom) the duplication starts with the same letter (C) as the letter that follows the duplication, in position 10. The consequence is a shorter break as the *k*-mer in position 3, that overlaps the duplication breakpoint exists in the reference. However, the duplication length is correctly computed as 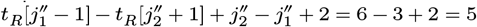.

We are finally able to identify duplications within a read in linear time. In order to limit the false positives, we introduce several strategies to mitigate this risk.

### 2.2 Mitigating false positives

False positive detection of duplication can be encountered because *k*-mers are short sequences that can occur by chance because of sequencing errors, or other events. We must not detect a duplication whenever a (few) short *k*-mers occur in the wrong place.

#### 2.2.1 Disambiguating positions

In the read trace, some *k*-mers may occur at several locations in the reference sequence. Of course only one (at most) is correct, thus only one is kept. Assuming we want to disambiguate *t*_*R*_[*i*] (*ie*. |*t*_*R*_[*i*]| > 1), we follow those rules:

1. First, if *t*_*R*_[*i* − 1] is non-ambiguous and *t*_*R*_[*i* − 1] + 1 appears in *t*_*R*_[*i*], then we retain this value.
2. Otherwise, among the possible positions in *t*_*R*_[*i*] we choose the one which minimises the distances (*ie*. the absolute values of the subtractions) with the neighbouring positions in *t*_*R*_ (both the original one and the one currently cleaned). The neighbouring positions considered are all the ones that don’t imply crossing a break, or a break can be crossed only if position *i* is just before or just after a break.

#### 2.2.2 Eliminating substitutions

Substitutions are the most frequent mutations and sequencing errors (at least for Illumina sequencers, see [24]). Therefore, many breaks will be due to substitutions. To limit the number of breaks to consider, and to have traces with more valid positions, we remove substitutions by correcting them (either mutations or sequencing errors). This also allows detecting duplications in spite of proximate substitutions, and it will improve the estimation of the quantification.

The correction is carried out as follows: let a break (*j*_1_, *j*_2_), we extract the *k*-mer starting at position *j*_1_. Since this *k*-mer doesn’t exist in *T*, as position *j*_1_ is the first one of a break, we try the three other nucleotides at position *k* − 1 in this *k*-mer. If one of the corrected *k*-mer occurs at a position *p* such that *p* − 1 occurs in *t*_*R*_[*j*_1_ − 1], then we correct the *k*-mer this way, and we move on to the next *k*-mer. In case the correction did not work, the *k*-mers in the break are not corrected.

#### 2.2.3 Simulating larger k-mers

Before reporting a duplication event, we make sure this event is robust by checking that *δ* consecutive positions in the trace are consistent. Verifying that *δ* consecutive positions are consistent is equivalent to considering *k* + *δ*-mers instead of *k*-mers. Thus, if we have a candidate break (*j*_1_, *j*_2_), we report this as an event if and only if *t*_*R*_[*j*_1_ − 1 − *i*] + *i* = *t*_*R*_[*j*_1_ − 1], for 1 ≤ *i* ≤ *δ*, and conversely iff *t*_*R*_[*j*_2_ + 1 + *i*] − *i* = *t*_*R*_[*j*_2_ + 1], for 1 ≤ *i* ≤ *δ. δ* is a user-defined parameter that is set to 2 by default.

Moreover, if two breaks (*j*_1_, *j*_2_) and 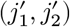 are consecutive by less than *δ* positions (*ie*. 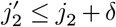), we merge them as soon as the indels they correspond to are longer than the one called using the merged break 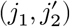.

### 2.3 Estimating duplication abundance

Identifying a duplication is not sufficient, as its abundance is also a prognosis factor. Rather than a raw abundance, a relative abundance is much more meaningful. We express the relative abundance as a VAF (Variant Allele Frequency) or an AR (allelic ratio). All the duplications sharing the same characteristics (same length and same starting and ending positions) are pooled together.

Briefly speaking, the abundance is computed from the number of reads in which a given duplication is found and the number of reads corresponding to the wildtype sequence. The read counts is further corrected to take into account the duplications that cannot be observed when they occur at one end of the reads.

More precisely, to determine the VAF or the AR we need to know, at the position of the duplication, both the coverage of the wildtype version and of the duplicated version, *ie*. the number of reads that cover the position of the duplication in the reference sequence and in the duplicated version. To do so, we use the disambiguated traces. The traces provide the positions that were covered in the reference sequence, we can thus count the coverage at each position in the reference. More formally, when *t*_*R*_[*i*] = *j*, we increment the coverage at position *j* in the reference sequence. This corresponds to the total coverage, which thus includes the coverage of the wildtype as well as the coverage of all the duplications or indels detected. We deduce the wildtype coverage by removing all the coverages from the duplications/indels that have been detected.

However, with our approach we cannot detect duplications that would occur at the start or at the end of a read. As explained previously, for a duplication to be detected, we need to have *δ* positions in the trace before and after the break. For a duplication creating a break of size *b*, we will need to have *t* = 2*δ* + *b* + *k* − 1 nucleotides in the read. That means that we will not be able to detect duplications occurring in the first *t* − 1 or the last *t* − 1 nucleotides of the reads. Thus, when quantifying a duplication, the quantification will be under-estimated because 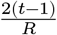 of the duplications could not be detected and, thus, counted, where *R* is the size of the reads. The counts are corrected to take this observation into account, and by assuming that any indel has a uniform probability of occurrence within the read. Let *q* be the raw quantification, *ie*. the number of reads in which the duplication was identified, the corrected quantification is 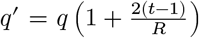. The VAF or AR are then computed using the corrected counts and the wildtype coverage at the position of the duplication.

### 2.4 Optimising data processing

As specified previously, we only search duplications in reads having enough *k*-mers coming from the reference sequence. However, in most cases most of the reads would not come from the region of interest. Thus, the most time-consuming step in our algorithm is to determine whether the read is coming from that region. To lower this time consumption, we use a heuristic: *k*-mers from the region of interest are stored in a Bloom filter [25].

The first step is therefore to query the Bloom filter to determine how many *k*-mers come from the region of interest in the read. If this value is above a threshold then the read goes to the following stages of the algorithm, otherwise it is discarded. By default, this threshold is set at 30 %.

Bloom filters do not yield any false negative, therefore we have the guarantee that this heuristic will not prevent us from identifying a read that actually comes from the region of interest. As the region of interest, namely exons 14 to 15 from the *FLT3* gene, is quite short the Bloom filter can be very small and will likely be stored in the CPU cache, which will allow very quick accesses.

### 2.5 Implementation and benchmarking

FiLT3r is implemented in C++ using the GATB library [26] and is freely distributed gitlab.univ-lille.fr/filt3r/filt3r.

We compared FiLT3r with state-of-the-art approaches. We made a preliminary assessment on our cohort with several other software detecting FLT3-ITD-ext, such as pindel [18], ITDSeek [27], ScanITD [28], getITD [29] and Genomon ITDetector [30] (see Supp Table 1). However, with the later publication of [19] we preferred referring to their independent assessment and chose to use in our paper the best tool they assessed: FLT3-ITD-ext [31]. We also added km [21] as it was not evaluated in Yuan et al’s benchmark and showed good performances in our preliminary assessment. km is an alignment-free approach already presented in the introduction. We also add that km is not focused on *FLT3* and can more generally detect variations in raw reads. *FLT3*-ITD-ext relies on a general-purpose aligner (BWA-MEM, [32]) to align reads to the reference sequence (the *FLT3* genomic sequence of exons 14-15).

**Table 1.**
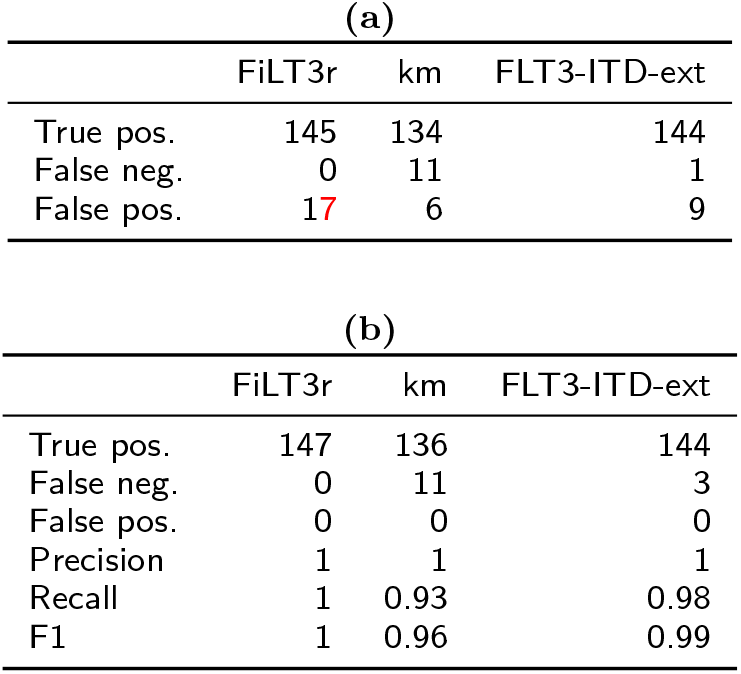
(a) Raw results of the three software packages assessed on the samples of 185 patients. Any result from a software is considered as a false positive as long as its quantification is at least 1% and it is not detected by the reference method. (b) Corrected results, after taking into account two Sanger sequencing to verify the two *FLT3*-ITDs detected by two software, and by considering as false positive only *FLT3*-ITDs detected above 1% by a software in any negative sample.

| (a) |  |  |  |
| --- | --- | --- | --- |
|  | FiLT3r | km | FLT3-ITD-ext |
| True pos. | 145 | 134 | 144 |
| False neg. | 0 | 11 | 1 |
| False pos. | 17 | 6 | 9 |

**Table 1** (a) Raw results of the three software packages assessed on the samples of 185 patients.
| (b) |  |  |  |
| --- | --- | --- | --- |
|  | FiLT3r | km | FLT3-ITD-ext |
| True pos. | 147 | 136 | 144 |
| False neg. | 0 | 11 | 3 |
| False pos. | 0 | 0 | 0 |
| Precision | 1 | 1 | 1 |
| Recall | 1 | 0.93 | 0.98 |
| F1 | 1 | 0.96 | 0.99 |

FiLT3r was launched with a *k*-mer size of 12^[1]^, km with a *k*-mer size of 31 (as advised by km authors) and with a threshold for duplication detections of .01 (thresholds .1 and .001 were attempted, but the best results were achieved with a threshold of .01), Jellyfish parameters were identical to the ones used in km publication (`-C -L 2 -Q+ -s 1000000`). FLT3-ITD-ext was launched with default parameters. For FLT3-ITD-ext, the deletions were not considered as it would have lead to too many false positives. Within a sample, all the duplications of the same length are pooled together to take into account the fact that the allelic ratio method only differentiates duplications on their lengths. The experiments can be reproduced with a Snakefile available on the Git repository of the software.

All the experiments were performed on a server with 24 Intel® Xeon® CPU E5-2420 and 193 GB of RAM. The data was stored on a NAS connected to the server through NFS. The programs were run on a single thread. User times and peak memory consumptions were measured.

### 2.6 Benchmarking data

We assessed the three software packages on high-throughput sequencing data of 185 patients aged from 18 to 60 years old, diagnosed with *de novo* or secondary AML at the haematology laboratory of the University Hospital of Lille. This population was composed of 114 *FLT3*-ITD positive patients and 71 *FLT3*-ITD negative patients, according to the gold-standard fragment analysis [33]. The study was conducted according to the Declaration of Helsinki and was approved (nos. CSTMT289) by the Human Research Committee of Lille and the institutional review board of the Lille University Hospital Tumor Bank (certification NF 96900-2014/65453-1).

Exons 14-15 of *FLT3* were sequenced in all patients using the following methodology: the library was prepared by a capture method, targeting 62 genes with SureSelect-QXT (AGILENT®) according to the manufacturer’s protocol. Samples were sequenced with NextSeq Illumina® (2 × 121 bp). The region of interest was sequenced with a mean depth of 2116x (range: 421.32 – 4538.84). The base calling was performed with `bcl2fastq2` (v:2.20.0) and fastq were trimmed with `fastp` (v:0.20.0). The number of paired-end reads per sample ranged from 1.8 million to 35 million (median: 4.8 million). The sequences were deposited on SRA (see supplementary file 3). The quantification of the expected ITDs were obtained using the reference method (by capillary electrophoresis) explained in the introduction.

To assess how the three software behaves on datasets with very different characteristics, we used public RNA-seq data. RNA-seq is gaining importance in clinical settings and RNA-seq data differs widely from capture sequencing in terms of quantity of data, variation of expression and random starting positions of the reads.

We launched the three software on data from CCLE (Cancel Cell Line Encyclopedia) [34]. We randomly selected 76 samples from haematopoeitic and lymphoid tissues and 76 samples from lung tissues (list of accessions in Supplementary file 10). It is not known which samples should contain *FLT3*-ITD. However, from a biological point of view, we do not expect to detect any *FLT3*-ITD in lung tissues while we could expect some in haematopoeitic and lymphoid tissues. This dataset will help assess how the three software scale on massive datasets (from 112,328 reads to 1,364,951,510 reads, median: 72.9M) with an average read length ranging from 68nt to 150nt (median: 101nt).

Finally, to check that our results holds on controlled data, we simulated ITDs at different ratios with different qualities.

## 3 Results

### 3.1 Capture sequencing of 185 AML patients

The software results on HTS data were compared to DNA fragment analysis, the gold-standard method. Any duplication found by a software, at whatever abundance, and identified with the reference method was considered as a true positive. Any duplication found by a software with an allelic ratio (AR) above .01 and that was not identified by the reference method was for now considered as a false positive. After analysing the results in details, we will change this definition of false positivity as HTS appear to be more sensitive to detect lowly expressed ITDs.

Raw results are shown in table 1 (a) (detailed results for each duplication are shown in Supplementary file 2). FiLT3r and FLT3-ITD-ext showed similar performances. FiLT3r was slightly better in terms of true positives, but at the apparent cost of more false positives.

However, some (7/17) of the false positives were shared with either km or FLT3-ITD-ext. We repeated a fragment analysis on the two samples where the false positives were the most abundant, to assess whether they were real false positives. We eventually obtained a sequence by Sanger for both (see supplementary files 8 and 9). Two other duplications classified as false positives detected by FiLT3r in SRR15006540 and undetected by the other software had cumulative lengths (16 and 56) corresponding to the length of a duplication detected by the conventional method (72). We thus believe that those two duplications are not false positives and are a consequence of our more fine-grained method that is able to distinguish multiple duplications. This is illustrated in supplementary file 1, figure 1, with a dotplot of a read from SRR15006540 against our reference sequence. Finally, seven other false positives were due to short single nucleotide deletions that could easily be filtered out.

It is noteworthy that all the false positives were identified in positive samples (apart from one of the Sanger-confirmed duplication) and at lower ratios than the duplication identified by the reference method. Thus, we consider that our original definition of *false positive* was too broad. Instead, we switch to the following, more strict definition: a false positive corresponds to any duplication detected at the threshold of .01 in a negative sample. No software detected a duplication above that threshold among the 71 negative samples. Hence, the most important metric in this context is the false negativity. We show a corrected version of the results in table 1 (b) where false positivity is thus at 0 and where the two Sanger-confirmed sequences are moved from the false positives to the true positives for FiLT3r and km (and added to the false negatives for FLT3-ITD-ext). Among those 185 samples, FiLT3r displayed perfect results as it did not report any false positive nor false negative result. FLT3-ITD-ext discarded all duplications whose length is not a multiple of 3. In AML, it was thought that the *FLT3*-ITD duplication should have a length multiple of 3. However, in our cohort, we showed that two duplications that had not been identified with the reference method did not have lengths multiple of 3. For km the false negatives are of varying lengths (from 6 nt to 126 nt) but with low concentrations. All the false negatives but one has a quantification ≤ 0.03 with the reference method, while the first half of the most abundant duplications were correctly called by km. Also, km correctly detects 10 duplications but is unable to provide a reliable quantification (it gives zero instead).

Beyond the correct detection of ITDs, it is important to also correctly assess the quantification of those duplications as they are meaningful for the prognosis of the patients. Fig. 4 shows the quantification computed by the three software packages for the duplications found. Their quantifications are compared to the one found by the reference method. With FiLT3r and FLT3-ITD-ext the quantification is closer to the fragment analysis quantification (respectively *r* = .93 and *r* = .88 for the log-transformed quantifications), compared to km (*r* = .76). One underestimation of FiLT3r quantification corresponded to the 72nt duplication we previously detailed. Interestingly, the other duplication FiLT3r largely underestimated (in SRR15006376) was also underestimated by a factor of 10 by FLT3-ITD-Ext and was not detected by km.

**Figure 4.**
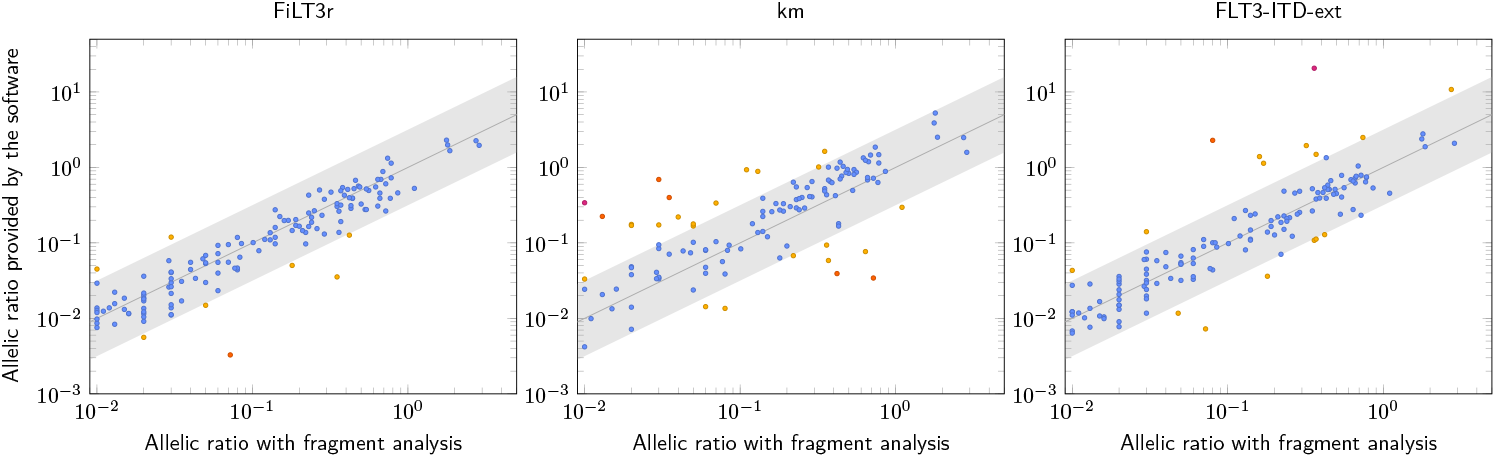
Quantification of duplications found with FiLT3r, km (threshold .01) and FLT3-ITD-ext compared with the fragment analysis method. The grey straight line corresponds to *y* = *x* and ideally the dots should be aligned along that line. The grey area is centred around that line and its width is of one log. The hotter the colour of the dots, the higher the quantification error. Only 8 results are not within this grey area for FiLT3r, 27 for km and 16 for FLT3-ITD-ext.

Regarding time and space consumption, FiLT3r showed the best performances (see Fig. 5 and 6). We also show the time and memory usage for counting *k*-mers, with Jellyfish, which is a required step for km.

**Figure 5.**
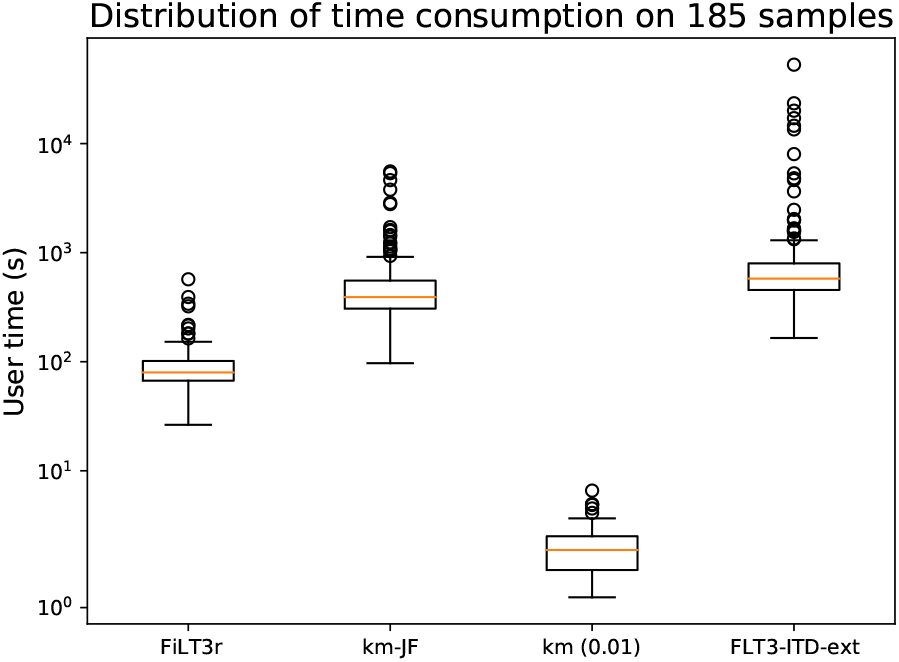
Time consumption of FiLT3r, km and FLT3-ITD-Ext on the 185 samples analysed. km-JF is the time taken by Jellyfish (preliminary step required for km), km (0.01) is the time taken by km with the corresponding threshold.

**Figure 6.**
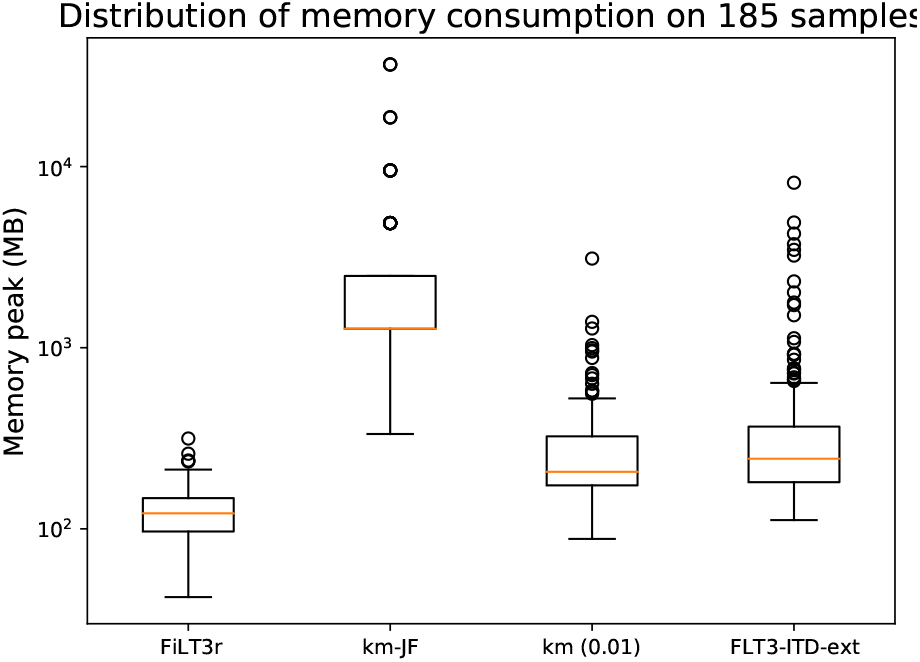
Memory consumption of FiLT3r, km and FLT3-ITD-Ext on the 185 samples analysed. km-JF is the memory taken by Jellyfish (preliminary step required for km), km (0.01) is the memory taken by km with the corresponding threshold.

In fig. 5 we see that km at thresholds 0.01 is very fast as it systematically took less than 10 seconds to detect the duplications. However, this does not include Jellyfish time, which must be launched on each sample. While Jellyfish is a very efficient *k*-mer counter, it was 2-3 orders of magnitude slower than km itself. FiLT3r’s median user time (65 s) is 6 times quicker than Jellyfish’s and 9 times quicker than FLT3-ITD-Ext’s. In some cases FLT3-ITD-Ext can be very time-consuming, taking several hours, while FiLT3r never took more than 10 minutes. The progression margin for FiLT3r is quite low as `gunzip` on the same files took about a third of the time took by FiLT3r (see supplementary figure 2).

For FiLT3r it appeared that using a Bloom filter dramatically speeded up the software as deactivating the Bloom filter makes FiLT3r one order of magnitude slower (see supplementary figure 3).

Memory consumption was very limited in FiLT3r as we stored a Bloom filter and a hash table of the reference sequence. We also stored reads that match the reference sequence in main memory, but there were generally just a few thousands of them. FLT3-ITD-Ext median memory consumption was close to FiLT3r’s as it was twice as space consuming. However, in some cases the space consumption could reach several GB while FiLT3r used at most 300 MB. Jellyfish memory usage was systematically above that highest memory usage recorded for FiLT3r. We could still improve FiLT3r memory usage by storing on the disk the reads matching the reference sequence. This would however lead to a time penalty.

### 3.2 152 RNA-Seq samples from CCLE

As expected none of the software detected a *FLT3*-ITD among the 76 lung samples. This tends to confirm the three software have a very low false positive rate. However on 10 samples (all with more than 1 billion reads), FLT3-ITD-ext had to be killed before the results were output as it took too much time (more than 24 h) or too much memory (more than 100 GB).

On the 76 haematological samples, some *FLT3*-ITD were detected by the three software. At the same 0.01 threshold as for the capture sequencing, FiLT3r detected 3 ITDs that were also detected by km, although one of them was only detected by km with `-p 0.001` not with `-p 0.01`. FLT3-ITD-Ext detected the remaining two.

km additionally detected a 60nt ITD in SRR8657348 that was not detected neither by FiLT3r nor FLT3-ITD-Ext. However this 60nt duplication seemed to be an artifact from a 30nt duplication that was actually detected by the three software. km seemed to detect a duplication of the duplication, however we found no read with evidence of such an event.

FLT3-ITD-ext detected three ITDs above the .01 threshold, two of which were not detected by FiLT3r or km: a 108nt ITD in SRR8615750, and a 105nt ITD in SRR8615696. The third ITD (a 78nt duplication in SRR8615696) was also detected by FiLT3r but at a much lower AR (.0007 instead of .02). In this sample we found evidence of only two reads containing the ITD breakpoint, casting doubt on the .02 AR given by FLT3-ITD-Ext. Moreover, km did not detect this ITD but was launched with options -p 0.01 or -p 0.001 and thus could not detect a very lowly expressed ITD. Regarding the 108nt ITD only found by FLT3-ITD-Ext, we found evidence of such a breakpoint in only one read. Concerning the 105nt ITD, the sequence reported by FLT3-ITD-Ext does not align in full-length on the reference, only the first 70nt do align. The last 35nt did not align on NCBI Blast non-redundant nucleotide sequences either. See supplementary file 1 for more details.

Overall, FiLT3r completed each task in less than 3 hours with less than 32 MB or RAM (median: 344 s and 26 MB). km (including Jellyfish step) took at most 9.5 hours and 37 GB of RAM for one sample (median: 962 s and 2.5 GB). FLT3-ITD-ext median time was 4400s with a median memory usage of 799 MB of RAM.

### 3.3 Simulated data

We simulated FLT3-ITDs using itdsim which is part of ITDseek [27]. Then the ITDs were sequenced *in silico* using `art` with the MiSeq v3 profile of errors [35]. The detailed results are presented in the supplementary material. In short, they confirmed the results obtained on real data. We also noticed that, the more sequencing errors there are, the more false positives FiLT3r has. However, those false positives are always quantified at ratio below .001. Contrarily to km or FLT3-ITD-ext, FiLT3r did not miss any ITD on the datasets with a normal coverage. FiLT3r also provided the best trade-off between quantification and F1 score.

## 4 Conclusions

We introduced FiLT3r, a time and memory efficient algorithm implemented in an open-source C++ software, which showed very good detection and quantification performances compared to state-of-the-art software. On our capture dataset, FiLT3r had a perfect recall, which was not the case of the other software. Even the reference fragment analysis method, exhibited two false negatives, as shown by further Sanger sequencing of the samples. Similarly to the other software, FiLT3r had a perfect precision. FiLT3r performances were better than that of FLT3-ITD-ext, the best software identified so far by an independent benchmark, which is alignment-based. Moreover FLT3-ITD-ext could not run on several RNA-seq dataset due to large resource requirements (both time and memory). In two cases on RNA-seq dataset, FLT3-ITD-ext reported quantification that appeared largely overestimated.

This illustrates that alignment-free algorithms can be more efficient than alignment-based algorithms while using a fraction of the resources they need. Beyond the *FLT3*-ITD analysis, FiLT3r can also be used to detect duplications in any gene as soon as the reference sequence is known. We plan to apply our method to other genes in other contexts, as it has been shown that tandem duplications can be prognostic signatures in other cancers (such as gastric cancers [36]).

## Supporting information

Supplementary material (including supplementary figures)

Supplementary file 11 - Detailed results for the simulated data with 150bp reads

Supplementary file 12 - Detailed results for the simulated data with 250bp reads

Supplementary file 13 - Detailed results for the simulated data with 250bp reads high-quality reads

Supplementary file 12 - Detailed results for the simulated data with 250bp reads and a low coverage

Supplementary file 12 - Detailed results for the simulated data with 250bp low-quality reads

## Ethics approval and consent to participate

The study was conducted according to the Declaration of Helsinki and was approved (nos. CSTMT289) by the Human Research Committee of Lille and the institutional review board of the Lille University Hospital Tumor Bank (certification NF 96900-2014/65453-1). The patients gave their written informed consent.

## Availability of data and materials

The software is available on the Git repository at gitlab.univ-lille.fr/filt3r/filt3r The experiments made in this paper can be reproduced using the Snakefile in the `reproducibility/` directory of the aforementioned Git repository. The data analysed in the publication is publicly available on SRA. The accessions of the datasets are provided in the Supplementary files.

## Consent for publication

Not applicable

## Authors’ contributions

AB, SD, MF and MS implemented the software. AB and MS performed the benchmarks. MS supervised the algorithmic design. ND, SG, MB, MD, RJ, LF, ON, CP performed the molecular studies. KCL ensured the data management. LG, RI, HD and MH provided samples and clinical data. HD and MH were the clinical investigators of the study. CP coordinated the biological collection. AB and MS wrote the manuscript. ND, MF, MD, RJ, RI corrected the proofs. All the authors approved the manuscript.

## Acknowledgements

We are grateful to É. Audemard for valuable discussions on their publications and software. We are also grateful to the bioinformatics service in CHU Lille for their helpful advices.

## Footnotes

[1] For a discussion on the choice of *k*, see the Supplementary Material, section *Choosing the parameters*.

