## Supplementary material (including supplementary figures) for "Frugal alignment-free identification of FLT3-internal tandem duplications with FiLT3r"

#### Contents

**1 Preliminary assessment of software to find *FLT3*-ITDs 2**

**2 Dotplot of a read with two duplications 2**

**3 Extra time consumption plots of FiLT3r 2**

**4 Choosing the parameters 2**

**5 Downsampling 6**

**6 ITDs detected by a single software from the CCLE dataset 7**

### 1 Preliminary assessment of software to find *FLT3*-ITDs

| | FiLT3r | Km | FLT3-ITD-ext | Pindel | ITDSeek | ScanITD | getITD | GID |
|---|---|---|---|---|---|---|---|---|
| True pos. | 147 | 125 | 144 | 123 | 50 | 120 | 123 | 115 |
| False neg. | 0 | 22 | 3 | 24 | 97 | 27 | 24 | 32 |
| False pos. | 0 | 0 | 0 | 0 | 5 | 1 | 0 | 2 |
| Precision | 1 | 1 | 1 | 1 | 0.91 | 0.99 | 1 | 0.98 |
| Recall | 1 | 0.85 | 0.98 | 0.84 | 0.34 | 0.82 | 0.84 | 0.78 |
| F1 | 1 | 0.92 | 0.99 | 0.91 | 0.50 | 0.89 | 0.91 | 0.87 |
| $r$ | 0.93 | 0.75 | 0.88 | 0.91 | 0.5 | 0.57 | 0.8 | 0.84 |

Table 1: Preliminary assessment made before the publication of Yuan et al (2021) on our cohort of 185 patients. Some results may differ from what is shown in the main article as the parameters were not necessarily the same.

### 2 Dotplot of a read with two duplications

See fig. 1.

### 3 Extra time consumption plots of FiLT3r

See fig. 2 for the comparison between FiLT3r and gunzip and fig. 3 to see the impact of the Bloom filter on FilT3r's time consumption.

### 4 Choosing the parameters

The value of the $k$-mer chosen for FiLT3r should be long enough to be specific but also short enough to ensure a high sensitivity.

We show in the supplementary file 4, experiments with values of $k$ ranging from 8 to 15. We obtain the same results as the ones presented in the main

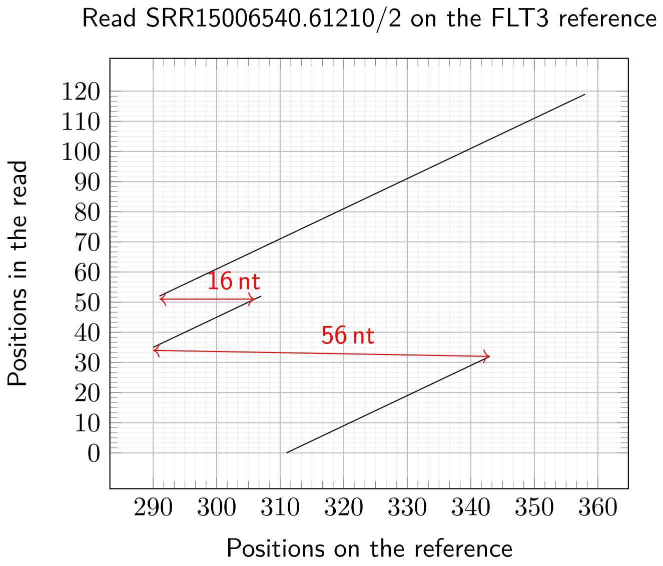

Figure 1: Dotplot of read SRR15006540.3329410/2 against the reference sequence used by FiLT3r. We see that the alignment is split in three different pieces, corresponding to two distinct duplications. The read starts aligning at position 311 in the reference and continues until read position 32 which aligns at position 343 in the reference. Then the alignment continues at position 35 in the read which aligns at position 290 in the reference, the alignement continues until position 52 in the read which aligns at position 307 in the reference. Finally another alignment starts at position 52 in the read on position 291 of the reference and continues until the end of the read. The red arrows indicate the duplication lengths.

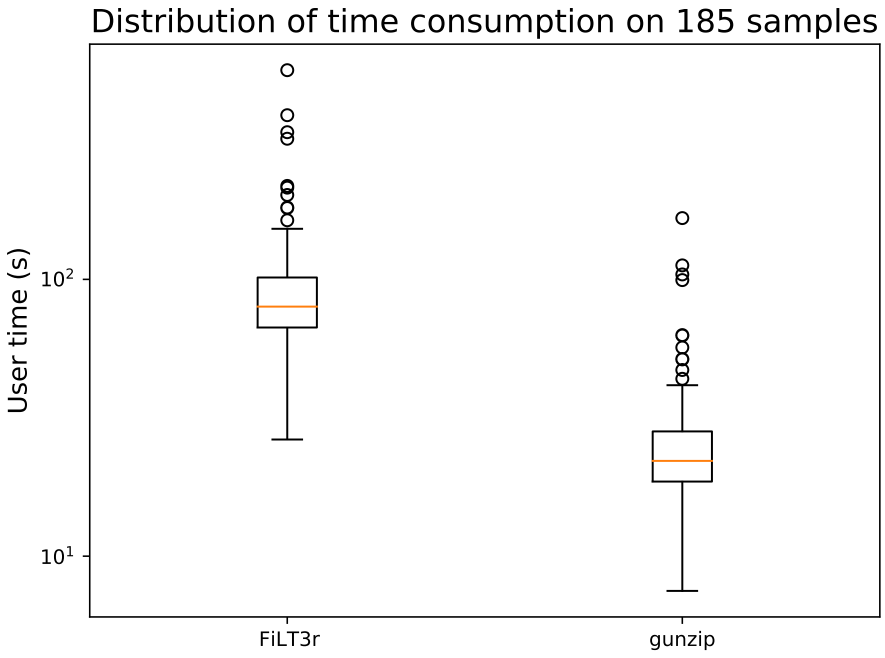

Figure 2: Time consumption comparison for FiLT3r and gunzip.

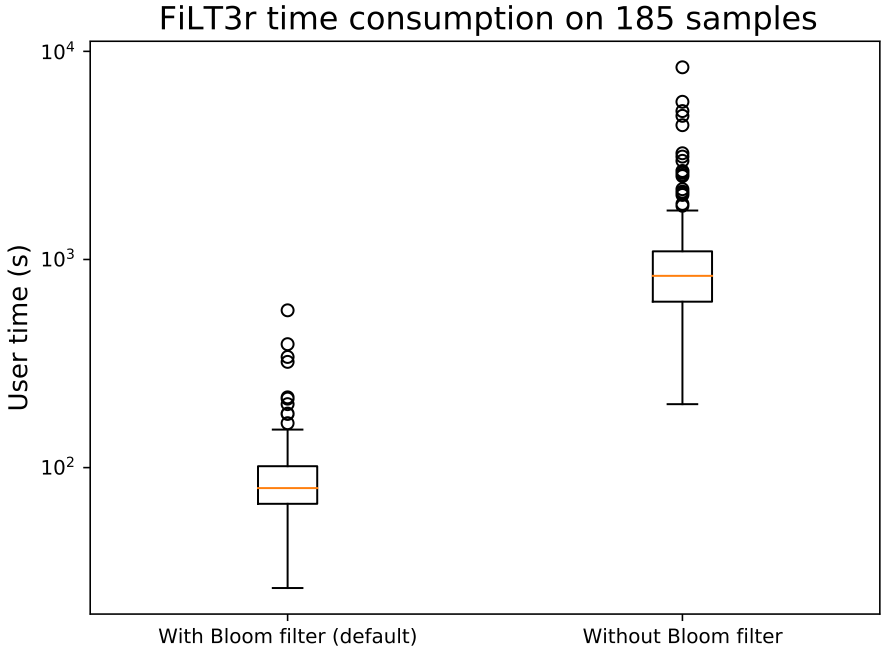

Figure 3: Time consumption of FiLT3r with and without Bloom filter. FiLT3r is much quicker when using a Bloom filter (which is the default). The median user time with a Bloom filter was 65 s and 834 s without.

article with $k = 11$ to 15. With $k = 10$, there is one false negative. The F1 score is still better than either km of FLT3-ITD-ext. The false negative is due to a redundancy within the $k$-mers which “hides” a break corresponding to a duplication.

At $k = 8$ the number of false negatives remain low (4) but the number of false positives is much larger (73). Those false positives are due to spurious hits of $k$-mers, as $k$ is too short. Actually the value of $k$ should be changed together with the threshold that determines the percentage of $k$-mer that must match the reference sequence as well as with the $\delta$ value used to simulate longer $k$-mers. In the supplementary file 5, we show experiments at $k = 8$ where the number of false positives is brought back to 0, with “only” 4 false negatives. Such results are thus similar to FLT3-ITD-ext, however the quantification we obtain with this $k$ is worse as the correlation coefficient with the reference method is about .8 instead of .93 with $k = 12$.

We also performed experiments with varying thresholds for the percentage of $k$-mers that must match the reference within a read so that it can be further analysed. We obtained the same results for percentages ranging from 20 % to 70 %. With higher percentages ($\geq$ 80%), there are many false negatives as the threshold becomes too stringent. On the contrary, at 10 %, the threshold is too permissive and we have a few false positives. The detailed results are provided in supplementary file 6.

### 5 Downsampling

For each sample we generated a file containing one tenth and one percent of the original reads. The paired reads that were kept were randomly chosen.

We emphasize that we did not expect the three software to be able to still report all the duplications as the initial coverage was chosen to be able to reach a 1% allelic ratio. As on average the coverage was 2116x, sampling one read out of ten, we can reach at most a .5% detection threshold (*ie.* a single read contributes to .5% of the coverage). However some samples have a lower coverage (min: 421x). When sampling one read out of a hundred, we can reach at most a 5% detection threshold, meaning that many duplications would be lost as a third of them were detected below that threshold with the gold-standard method.

We indicate in the table below how many duplications were lost when sampling 10 % or 1 % of the reads, compared to the results on the whole dataset. Note that for FiLT3r we set the parameter `--min-nb-duplication` to 1 (default is 2) as FLT3-ITD-ext reports duplications that are detected in a single read and for `km` we set `-L 1` for `jellyfish count` and `-c 1` for `km find_mutation`

| | FiLT3r | Km | FLT3-ITD-ext |
|---|---|---|---|
| Duplications lost with 10% of the reads | 3 | 5 | 5 |
| Duplications lost with 1% of the reads | 49 | 53 | 49 |

The 3 duplications lost by FiLT3r are also lost by FLT3-ITD-ext and, for one of them, was already not detected by km in the full dataset. Those duplications are:

- the 57nt duplication in SRR15006388, with an allelic ratio of 1% according to the reference method
- the 24nt duplication in SRR15006460, with an allelic ratio of 3% according to the reference method
- the 111nt duplication in SRR15006521, with an allelic ratio of 3% according to the reference method

Thus all the software seem to be similarly impacted by the downsampling.

However we should also note that FiLT3r seems to have much more false positives (32) in the 10% subsample. Actually this is due to 1 nucleotide deletions. For the sake of comparison with the reference method, which is not sequence nor position-based, we gather all the insertion/deletion/duplication events of the same length in a single one. Therefore all 1-nt deletion will be gathered in a single one which, by chance, may be enough to reach the 1% threshold in a subsample of the real data. 29 of the 32 false positives are such cases, that are therefore artifacts of how we compare results and that could easily be discarded if one wanted. Note that, as for the main results, we do not report the deletions of FLT3-ITD-ext. The remaining three false positives are:

- a 3-nt duplication in SRR15006427, which is also detected by FLT3-ITD-ext, in the same subsample
- a 6-nt deletion in SRR15006449, which is also detected by km in the same subsample.
- a 57-nt duplication in SRR15006457, which is also detected by km in the same subsample.

The details of the results are provided in the supplementary file 7.

Surprisingly the correlation of the quantifications with the reference method are little affected by the 10% subsampling.

The results in the 1% subsamples are poor for the three software but this was expected as the coverage was not sufficient to reach the quantification threshold.

### 6 ITDs detected by a single software from the CCLE dataset

#### 6.1 60nt ITD in SRR8657348

This ITD was only detected by km. According to km output the ITD is ACGTTGATTTCAGAGAATATGAATATGATCACGTTGATTTCAGAGAATATGAATATGATC, which is a duplication itself of the 30nt ACGTTGATTTCAGAGAATATGAATATGATC. This 30nt ITD was detected by the three software.

We searched all the reads containing AATATGAATATGATCACGTTGATTTCA-GAG (ie. the last 15nt of the duplication followed by the first 15nt of the duplication) to identify reads containing the duplication. If there was a duplication of the duplication, we would expect the ITD to be followed or preceded by another occurrence of the ITD.

Aligning all those reads gives the following consensus sequence:
CTCCTCAGATAATGAGTACTTCT
ACGTTGATTTCAGAGAATATGAATATGATC
ACGTTGATTTCAGAGAATATGAATATGATC
TCAAATGGGAGTTTCCAAGAGA

Thus we do not find any clue of the third occurrence of the 30nt ITD that would justify considering a 60nt ITD

##### 6.2 108nt ITD in SRR8615750

This ITD was only detected by FLT3-ITD-ext. The software reports the following ITD: AAATCAACGTAGAAGTACTCATTATCTGAGGAGCCGGTCAC-CTGTACCATCTGTAGCTGGCTTTCATACCTAAATTGCTTTTTGTACTTGT GACAAATTAGCAGGGTT. We searched within the reads for the last 10nt of this ITD, followed by the first 10nt of the ITD, to search for the breakpoint of the ITD. We found a single occurrence among the 153M paired-end reads. However FLT3-ITD-ext reported it with an AR of 4.7. We did not find any evidence of such a highly expressed 108nt ITD in this dataset.

##### 6.3 105nt ITD in SRR8615696

This ITD was only detected by FLT3-ITD-ext. The software reports the following ITD: CGTAGAAGTACTCATTATCTGAGGAGCCGGTCACCTGTACCATCT-GTAGCTGGCTTTCATACCTAAATTGCTTTTTGTACTTGTGACCGGCTCCTCTGAAATCAG

The sequence itself does not belong to the FLT3 exon14-15 locus: only the first 70nt align on the reference, the remaining 35 nt (CTTTTTGTACTTGT-GACCGGCTCCTCTGAAATCAG) were searched with BlastN on the non-redundant nucleotide collection, no hit was found.

#### 7 Simulated data

We simulated FLT3-ITDs using itdsim which is part of ITDseek [1]. Then the ITDs were sequenced *in silico* using `art` with the MiSeq v3 profile of errors [2]. ITDs were simulated at seven differing ratios, from .1 to .001, with 10 ITDs at each ratio, resulting in 70 ITDs. We simulated datasets with 150bp paired-end reads, others with 250bp paired-end reads. For the 250-bp datasets, we generated reads with normal, high and low qualities ((qualities 10 times better, respectively lower, than usual with the `art` MiSeq v3 error profile), we also generated a dataset with a low coverage (10 times lower than normal). Each ITD was simulated in an independent dataset, with 60,000 reads from the wildtype sequence for 150bp reads, 30,000 reads for the 250bp datasets (note that this is not the coverage but the number of reads covering the whole wildtype sequence).

For those simulated data, we analyzed all the ITDs detected above a ratio of .0005, which is much lower than on the real data to stress the software with small coverage of the ITDs. As expected, FLT3-ITD-Ext didn't detect duplication lengths that were not a multiple of 3. The results for FLT3-ITD-ext below are always restricted to the ITDs that are a multiple of 3, FLT3-ITD-Ext missed 3 of them.

#### 7.1 Results on 150bp reads

| | FiLT3r | Km (p=.01) | Km (p=.001) | FLT3-ITD-ext* |
|---|---|---|---|---|
| True pos. | 70 | 34 | 60 | 24 |
| False neg. | 0 | 36 | 10 | 3 |
| False pos. | 1 | 0 | 0 | 3 |
| F1 | 0.99 | 0.65 | 0.92 | 0.89 |
| Quantification ($r$) | 0.99 | 0.94 | 0.86 | 0.97 |

FiLT3r had one false positive: a 520bp deletion. This is an actual false positive due to redundancy of some $k$-mers. By looking at all the results on this sample with FiLT3r we also notice the "converse' ' event (with a 520bp insertion). This would help filtering such events and such a filtering could be integrated in future versions of FiLT3r.

#### 7.2 Results on 250bp reads

| | FiLT3r | Km (p=.01) | Km (p=.001) | FLT3-ITD-ext* |
|---|---|---|---|---|
| True pos. | 70 | 40 | 64 | 22 |
| False neg. | 0 | 30 | 6 | 5 |
| False pos. | 2 | 0 | 0 | 12 |
| F1 | 0.99 | 0.73 | 0.96 | 0.72 |
| Quantification ($r$) | 0.99 | 0.96 | 0.85 | 0.89 |

All FLT3-ITD-ext false positives were found at low ratios (betwen .0005 and .0006). The two FiLT3r false positives were also found at low ratios (.0005 and .0008) and were single nucleotide deletion, probably due to simulated sequencing errors.

When we restrict to duplication lengths that are a multiple of 3, FLT3-ITD-ext still missed 5 of them (out of 27).

On real data, all those false positives would not be a problem as the detection threshold is much higher.

#### 7.3 Results on 250bp low-quality reads

| | FiLT3r | Km (p=.01) | Km (p=.001) | FLT3-ITD-ext* |
|---|---|---|---|---|
| True pos. | 70 | 41 | 58 | 0 |
| False neg. | 0 | 29 | 12 | 70 |
| False pos. | 22 | 0 | 0 | 0 |
| F1 | 0.86 | 0.74 | 0.91 | NA |
| Quantification ($r$) | 0.99 | 0.74 | 0.91 | NA |

FiLT3r had more false positive for this dataset because of the large number of sequencing errors (due to the simulated low quality run). However 21 out of 22 of those false positives were at a lower ratio than the ITD that had to be found since all the false positives were quantified between .0005 and .0009. In spite of those sequencing errors, FiLT3r is still able to determine the ITD quantification accurately. For the first time, FiLT3r had not the best F1-score, km (with p=.001) had a better one, while still missing 12 ITDs. However it should be mentioned that km (with p=.001) was very slow, taking on average 40 min for files that had less than 70,000 reads. FiLT3r completed the job in 9s on average. With this level of error, FLT3-ITD-ext is not able to find any ITD anymore.

#### 7.4 Results on 250bp high-quality reads

| | FiLT3r | Km (p=.01) | Km (p=.001) | FLT3-ITD-ext* |
|---|---|---|---|---|
| True pos. | 70 | 40 | 68 | 20 |
| False neg. | 0 | 30 | 2 | 4 |
| False pos. | 4 | 0 | 0 | 0 |
| F1 | 0.97 | 0.57 | 0.99 | 0.91 |
| Quantification ($r$) | 0.99 | 0.95 | 0.88 | 0.99 |

As expected the results on this dataset are better, as there are fewer sequencing errors. The F1 scores are globally high and the quantifications are accurate. km (p=.001) has the best F1 score but it has the worst quantification, so the best trade-off seems to be the results obtained by FiLT3r. The four FiLT3r false positives were single nucleotide deletion at very low ratio (between .0005 and .0008).

#### 7.5 Results on 250-bp reads with low coverage

| | FiLT3r | Km (p=.01) | Km (p=.001) | FLT3-ITD-ext* |
|---|---|---|---|---|
| True pos. | 55 | 37 | 40 | 10 |
| False neg. | 15 | 33 | 30 | 10 |
| False pos. | 0 | 0 | 0 | 8 |
| F1 | 0.88 | 0.69 | 0.73 | 0.53 |
| Quantification ($r$) | 0.94 | 0.75 | 0.64 | 0.79 |

For FiLT3r, all the false negative had a theoretical quantification $\leq .002$. For km, all the false negatives were at theoretical quantification $\leq .01$.

##### 7.6 Discussion of the results on simulated data

Overall, FiLT3r systematically obtained the best correlation for the quantification of the ITDs and never had any false negative on the full datasets. Globally, apart on the worst datasets (bad quality of low coverage), it systematically obtained very high F1 scores ($\geq .97$). Globally, km (with p=.001) also had good F1 scores, without any false positive, but the quantifications were less accurate. The ITDs missed were among the least abundant for km (but lowering the `p` parameter dramatically increases the computation time) but FLT3-ITD-ext also missed some ITDs among the most abundant ones (for instance one ITD at a ratio of .1 in the 250bp dataset and in the 250bp high-quality reads). However the downside is that FiLT3r had a few false positives, always quantified below .0009, even with a high level of sequencing errors. It will be a future investigation to determine how some of those false positives could be filtered out. In several instances, there are two false positives consisting in an indel of length $n$ and another one of length $-n$. Such events are not expected and could be filtered out, by taking some precautions.

Those results confirm what we observed on the real datasets: FiLT3r is more sensitive, but can have some false positives (mainly single nucleotide deletions) at low concentration, and has a better quantification than its counterparts (even with a lower coverage).
