## Supplemental Data 7 for "Frugal alignment-free identification of FLT3-internal tandem duplications with FiLT3r"

Applied Biosystems

GeneMapper 4.1

ANALYSES

### SRR15006459

| Sample File | Sample Name | OS | SQ |
|---|---|---|---|

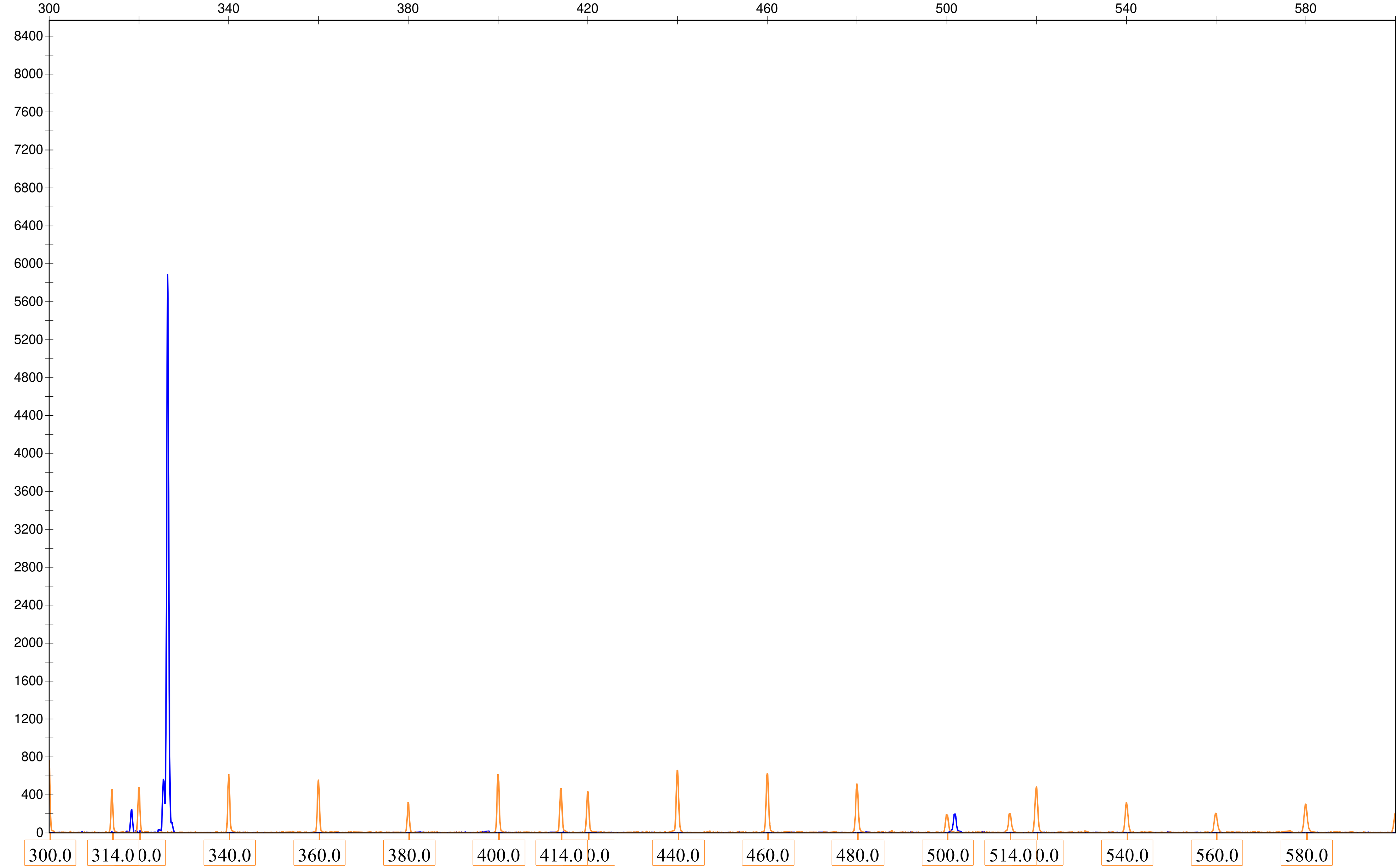

ANALYSES

SRR15006459

| | | Sample File Name | Size | Height | Area |
|---|---|---|---|---|---|
| 1 | ■ | [redacted] | 326.42 | 7772 | 61570 |
| 2 | ■ | [redacted] | 501.78 | 281 | 3139 |
