## Supplemental Data 8 for "Frugal alignment-free identification of FLT3-internal tandem duplications with FiLT3r"

Applied Biosystems
GeneMapper 4.1

### ANALYSES

#### SRR15006372

| Sample File | Sample Name | OS | SQ |
| --- | --- | --- | --- |

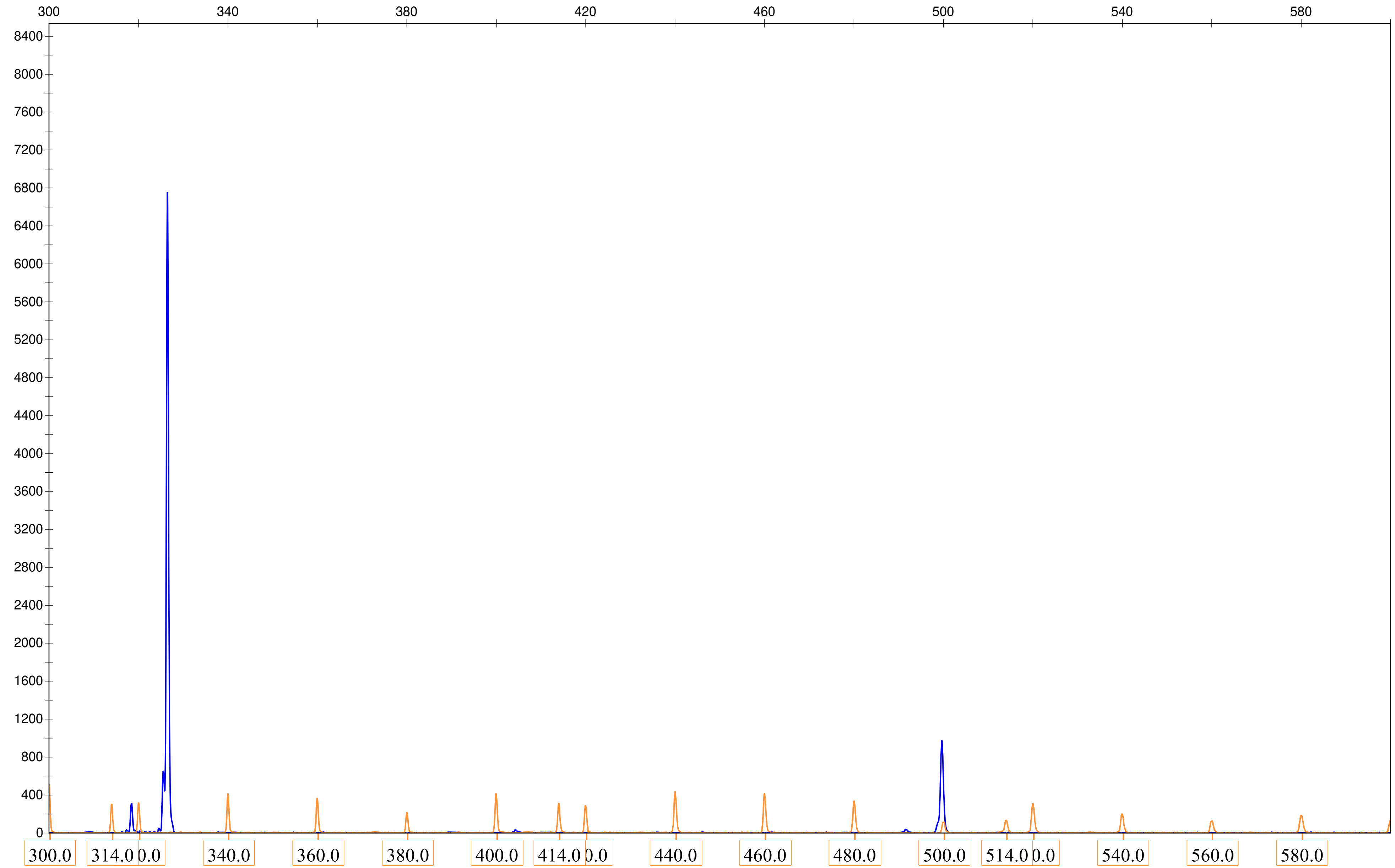

ANALYSES

SRR15006372

| | | Sample File Name | Size | Height | Area |
|---|---|---|---|---|---|
| **1** | | [redacted] | 326.49 | 6756 | 54376 |
| **2** | | [redacted] | 499.62 | 980 | 11292 |
